# A negative role for the interleukin-2-inducible T-cell kinase (ITK) in human Foxp3+ T_REG_ differentiation

**DOI:** 10.1101/386508

**Authors:** Polina Mamontov, Ryan A. Eberwine, Jackie Perrigoue, Anuk Das, Joshua R. Friedman, J. Rodrigo Mora

**Affiliations:** Janssen Research & Development, Spring House, PA 19477, USA

## Abstract

The Tec kinases ITK (interleukin-2-inducible T-cell kinase) and RLK (resting lymphocyte kinase) are critical components of the proximal TCR/CD3 signal transduction machinery, and data in mice suggest that ITK negatively regulates T_REG_ differentiation. However, whether Tec kinases modulate T_REG_ development and/or function in human T cells remains unknown. Using a novel self-delivery siRNA platform (sdRNA), we found that ITK knockdown in primary human naïve peripheral blood CD4 T cells increased Foxp3^+^ T_REG_ differentiation under both T_REG_ and T effector (Teff) cell priming conditions. ITK knockdown also enhanced the expression of the co-inhibitory receptor PD-1 on FoxP3+ T cells. T_REGS_ differentiated in vitro (iT_REG_) after ITK knockdown displayed suppressive capacity against effector CD4+ T cell proliferation. ITK knockdown decreased IL-17A production in T cells primed under Th17 conditions and increased Th1 differentiation. Finally, a dual ITK/RLK Tec kinase inhibitor blocked T_REG_ differentiation and T cell activation in general. Our data suggest that targeting ITK in human T cells may be an effective approach to boost T_REG_ in the context of autoimmune diseases, but non-specific inhibition of other Tec family kinases may broadly inhibit T cell activation.

## Introduction

Interleukin-2-inducible T-cell kinase (ITK) is a member of the TEC kinase family of non-receptor tyrosine kinases and mediates T cell signaling downstream of TCR activation [1]. Signaling through ITK modulates T cell activation, T helper cell differentiation, and thymic selection of developing thymocytes. ITK has been implicated as a critical node in T cell and NK cell mediated inflammation, leading to interest in developing therapeutics to modulate ITK function in autoimmune and inflammatory diseases [2, 3].

ITK is thought to drive Th2-mediated disease such as allergic asthma, and ITK^-/-^ mice exhibit significantly improved disease course and reduced bronchoconstriction after antigen re-challenge in ovalbumin sensitized mice [4, 5]. ITK has also been proposed to regulate the balance between inflammatory CD4+ Th17 cells and CD4+ regulatory T cells (T_REG_) in mice [6]. ITK is an important switch for Th1 and Th2 mediated immunity, and murine ITK deficiency results in reduced differentiation and effector cytokine production from Th1, Th2, and Th17 polarized CD4+ T cells while bolstering T_REG_ development [6-9].

However, since ITK is also involved in thymocyte development [10], studies in knock-out mice may not distinguish potential developmental defects in the immune system from the effects of ITK inhibition on the mature immune system. Although ITK also serves a scaffolding function for the docking of signaling intermediates (such as Vav1) that coordinate actin polymerization toward the TCR-APC interface [11], studies in kinase-dead ITK mutant mice have shown that ITK kinase activity was required for driving Th1, Th2, and Th17 differentiation [7, 8].

Resting lymphocyte kinase (RLK) is another member of the TEC family of non-receptor tyrosine kinases closely related to ITK. Less is known about RLK in T cell signaling and differentiation. While both ITK and RLK are activated by Src kinases downstream of the TCR signaling complex, RLK activation is independent of PI3K activation [12]. RLK is constitutively bound to the T cell plasma membrane via an N-terminal palmitoylation site, whereas ITK has a pleckstrin homology domain which requires PI3K-mediated PIP3 generation for recruitment to the plasma membrane after TCR activation [12-15]. Txk mRNA (encoding RLK) is also downregulated in T cells following activation [15]. Together this suggests that ITK and RLK play redundant as well as distinct roles in T cell development and activation. In fact, whereas ITK-/- mice exhibit impaired CD4+ and CD8+ T cell development, RLK deficiency alone does not affect T cell development. However, mice that are deficient in both ITK and RLK have a marked defect in T cell activation in response to anti-CD3, but not PMA, stimulation [1].

While ITK is required for IL-17a production in human T cell lines [14] and it also regulates Th17 and T_REG_ differentiation in mice [6], its role in human T_REG_ differentiation is not defined. Here we investigated the roles of ITK in human Foxp3+ T_REG_ differentiation and function using self-delivered siRNA (sdRNA) technology optimized to modulate ITK expression in resting primary human T cells. We found that ITK is a negative regulator of human T_REG_ differentiation under *in vitro* T_REG_, Th17, and Th1 polarizing conditions, and that ITK reciprocally regulates T_REG_ and Th17 differentiation from naïve human CD4+ T cells. ITK also regulates the expression of the co-inhibitory molecule PD-1 on *in vitro*-differentiated human FoxP3+ CD4+ T cells. Of note, pharmacologic inhibition of multiple TEC family kinases (ITK and RLK) blocked the T_REG_ bolstering effect of ITK knockdown, suggesting that RLK may be required for T_REG_ differentiation in the absence of ITK. Our data show for the first time that ITK regulates CD4+ T_REG_ and Th17 differentiation in primary human T cells and highlights the need for more specific pharmacologic targeting of TEC kinases in inflammatory and autoimmune disease.

## Results

### ITK knockdown in human naïve CD4 T cells promotes Foxp3^+^ T_REG_ differentiation

To assess the effect of ITK on human T_REG_ differentiation, we used a recently developed self-delivered siRNA technology (sdRNA; Advirna, Cambridge, MA) which circumvents the low efficiency of conventional gene knockdown methods in resting primary human T cells. ITK sdRNA treatment led to a 50%-80% reduction in ITK gene expression relative to a non-targeting sdRNA control (NTC) in human peripheral blood naïve CD4+ T cells (Figure 1A). CD4+ T cells were then differentiated under T_REG_, Th17, Th1, or non-polarizing conditions (Th0). Knockdown of ITK resulted in a 3-8-fold increase in the frequency of FoxP3+ T cells and a 2-5-fold increase in FoxP3 MFI under T_REG_, Th17 and Th1 polarizing conditions relative to NTC (Figure 1B, C). These results suggest an inverse relationship between ITK gene expression and FoxP3+ T cell differentiation in human CD4 T cells.

**Figure 1.**
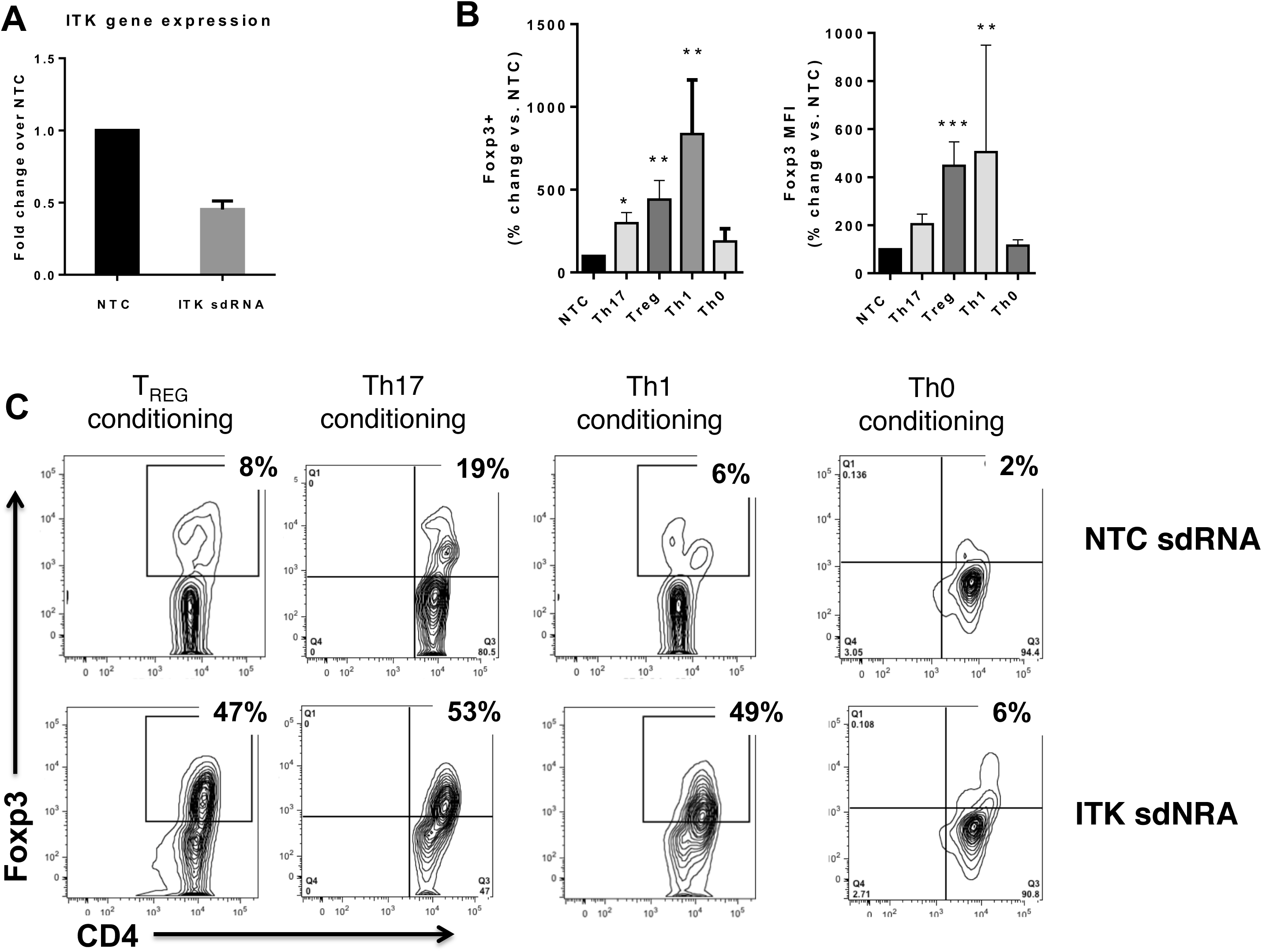
**A:** ITK mRNA expression in sdRNA treated cells was normalized by the ∆∆Ct method to the mean of endogenous control genes (TBP, IPO8, and HPRT1). ITK expression is shown as fold change relative to non-targeting sdRNA. N=5 donors; P<0.0001 (Wilcoxon Rank Test) **B and C**: Human naïve CD4+ T cells were treated with non-targeting (NTC) or ITK targeting sdRNA and then activated under Th17 (TGFβ, IL-2, IL-6, IL-23, IL-1β, anti-IFNγ, anti-IL-4), Th1 (IL-2, IL-12) or T_REG_ (IL-2, TGFβ) polarizing conditions. Four days later, CD4 T cells were analyzed for Foxp3 expression by flow cytometry. N=3-5 donors per condition, shown is mean +/-SEM. *p<0.05, **p<0.01. ***p<0.001 (ANOVA vs. NTC)

### ITK knockdown modulates Th17 and Th1 differentiation in human T cells

Deletion of *ITK* in mice leads to increased T_REG_ development under both T_REG_ and Th17 polarizing conditions [6]. While ITK is critical for driving Th2 mediated immune responses and effector cytokine production [9, 16, 17] some studies indicate that ITK is also required for Th1 effector cytokine production in mouse T cells [7, 9, 17].

Th17 cells produce the cytokine IL17A and express the chemokine receptor CCR6. We examined IL17A secretion and surface expression of CCR6 in human Th17 polarized cells upon ITK knock down and observed down-regulation of both markers relative to T cells treated with NTC (Figure 2A, B). Thus, analogous to mouse T cells [6], ITK reciprocally regulates the differentiation of T_REG_ and Th17 cells in human primary T cells.

**Figure 2.**
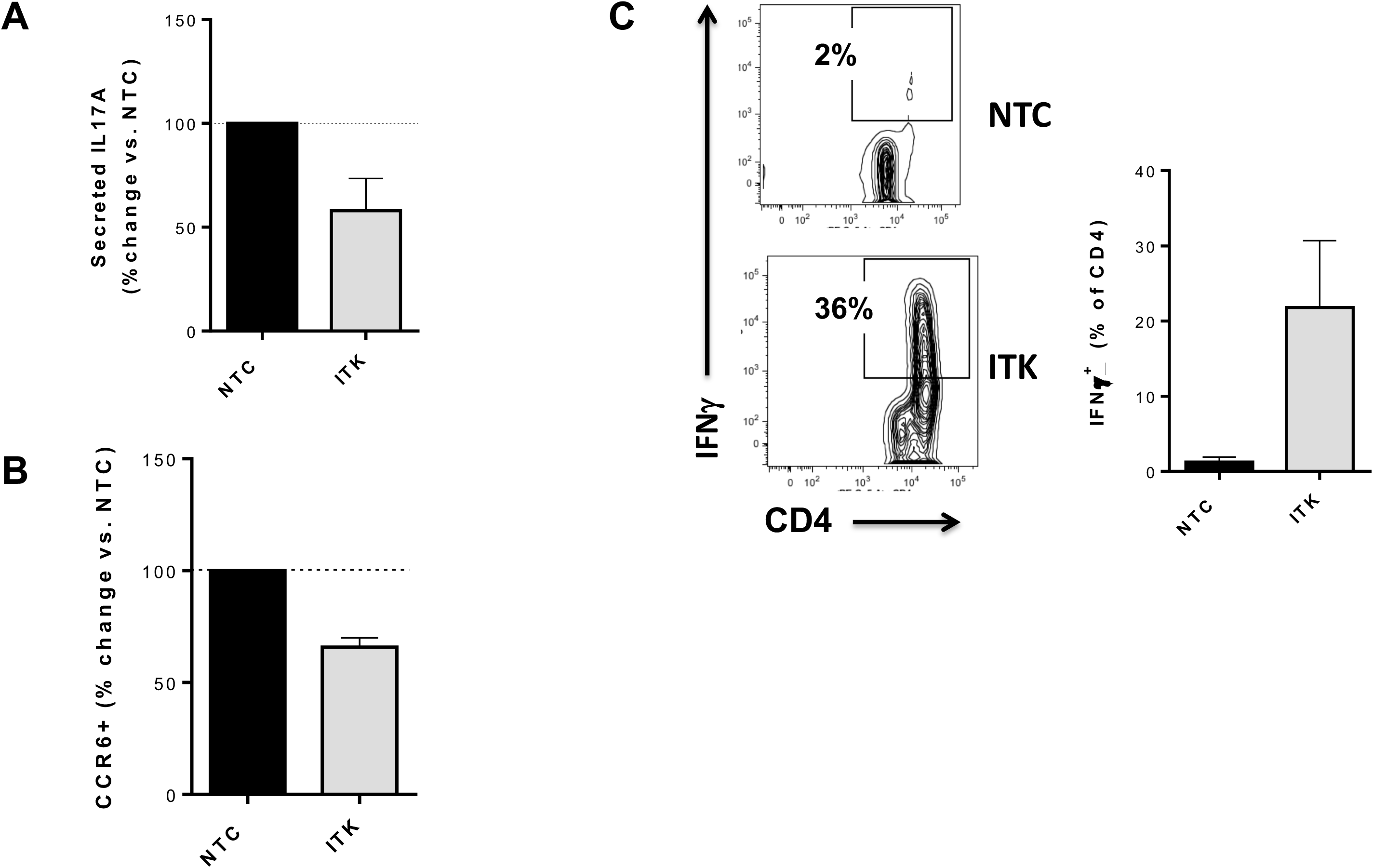
Human naïve CD4 T cells were treated with non-targeting (NTC) or ITK targeting sdRNA and then activated under Th17 (TGFβ, IL-2, IL-6, IL-23, IL-1β, anti-IFNγ, anti-IL-4) or Th1 (IL-2, IL-12) polarizing conditions. Four days later, culture supernatants were assessed for IL-17A by Luminex (**A;** N=5) and CD4+ T cells were analyzed for CCR6 (**B;** N=3) or INFɣ (**C;** N=3) by flow cytometry.

Although ITK knockdown led to increased development of FoxP3+ T cells under Th1 polarizing conditions, it also led to increased IFNγ expression (Figure 2C). This is in contrast to what has been observed in mice expressing a mutant form of ITK whose kinase activity can be selectively inhibited by small molecule inhibitors, where ITK kinase activity was required for Th1, Th2, and Th17 effector cytokine production [7], or in ITK-/- mice, which exhibit normal Th1 differentiation [5, 6].

### ITK knockdown increases the proportion of human T_REG_ expressing the co-inhibitory receptor PD-1

While FoxP3 is a requisite marker of bona-fide regulatory T cells, in humans it may not be sufficient to define functional T_REG_ cells. Thus, we further assessed surface expression of the co-inhibitory receptor PD-1 upon ITK knock down in T_REG_ polarized cells. Consistent with a negative role of ITK in functional human T_REG_ differentiation, ITK knockdown led to a significant increase in the proportion of FoxP3+ cells expressing the co-inhibitory molecule PD-1, suggesting that the increase in Foxp3+ T cells correlates with functional suppressive capacity (Figure 3).

**Figure 3.**
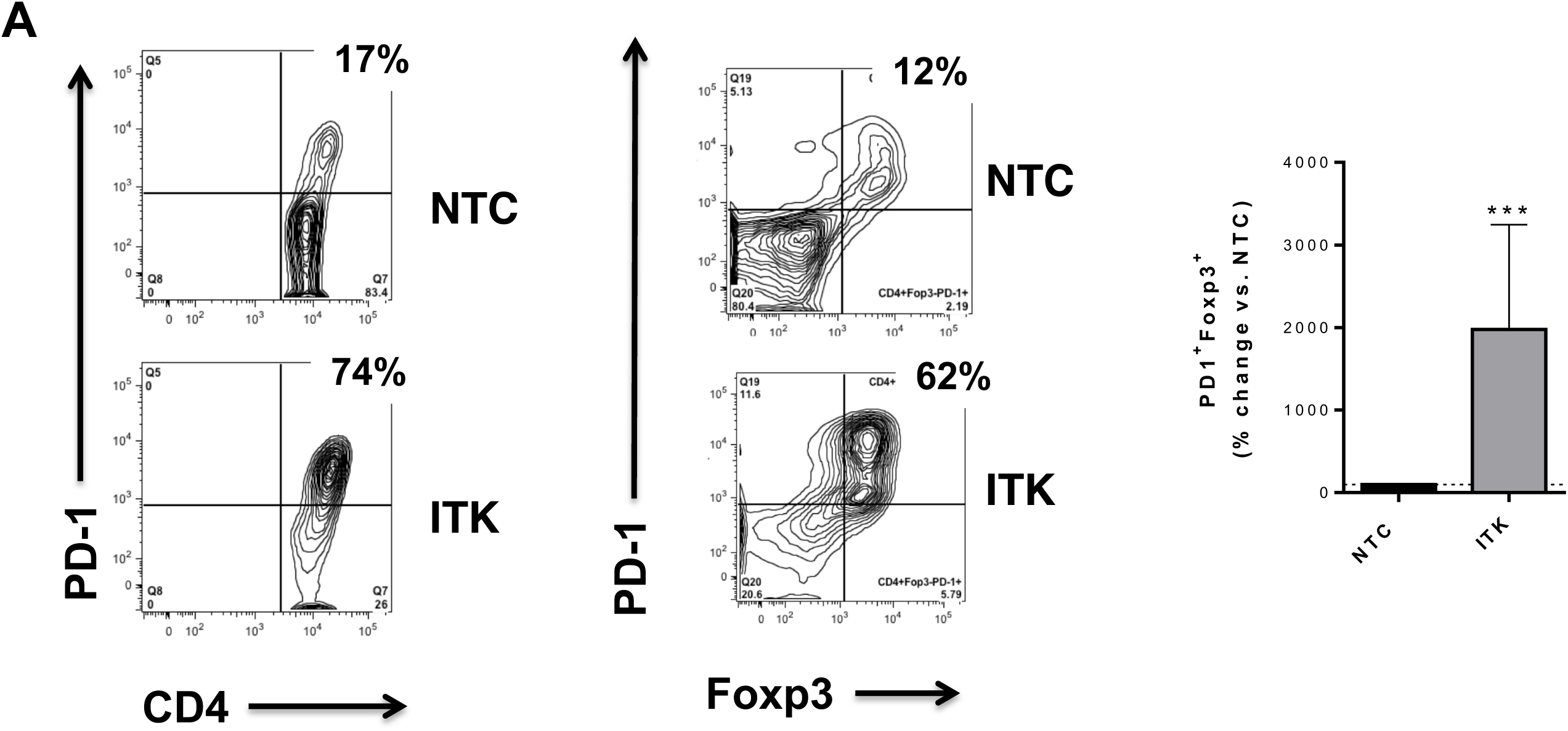
Human naïve CD4 T cells were treated with non-targeting (NTC) or ITK targeting sdRNA and then activated under T_REG_ (IL-2, TGFβ) polarizing conditions. Four days later the CD4 T cells were analyzed for Foxp3, and PD-1 expression by flow cytometry. N=7 donors. ***p<0.001 (Wilcoxon Rank Test)

### ITK knockdown increases the proportion of functionally suppressive human T_REG_

Given that ITK knockdown bolsters T_REG_ abundance and increases FoxP3 and PD-1 expression, we next examined the functionality of T_REGS_ differentiated *in vitro* in the presence of ITK sdRNA. *In vitro* differentiated T_REGS_ were co-cultured with CFSE-labeled CD4+ responder cells and anti-CD3 anti-CD28 activation beads, and responder T cell proliferation was measured based on CFSE dilution (Figure 4A).

**Figure 4.**
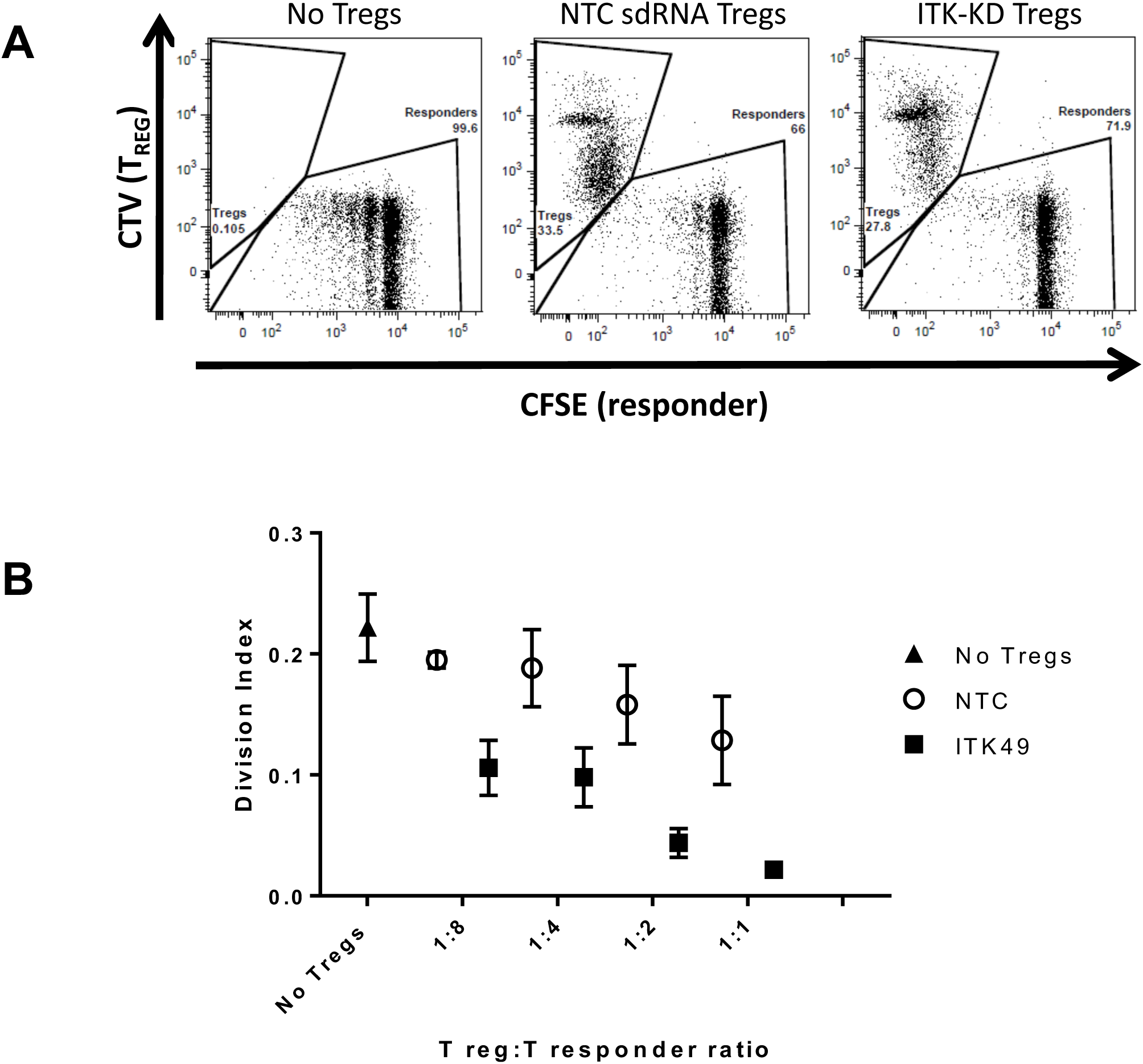
Human naïve T cells were treated with ITK or NTC sdRNAs and differentiated under T_REG_ polarizing conditions (iT_REG_). *In vitro* suppression assays were set up with iT_REG_ labeled with CellTrace Violet (CVT) and CD4+CD25-responder T cells labeled with CFSE in the presence of anti-CD3/CD28 activation beads. FACS analysis was performed after 4 days (**A**, 1:1 cell ratio shown) and the FlowJo proliferation platform was used to calculate the division index (DI; **B**). DI= the average number of cell divisions of the original population including undivided cells. N=4 donors.

T_REG_ differentiated *in vitro* in the presence of both ITK and NTC sdRNA were suppressive against effector cell proliferation in a T_REG_-dose dependent manner (Figure 4B). While T_REGS_ differentiated in the presence of ITK sdRNA appeared to exhibit enhanced suppressive capacity in this assay, ITK knockdown leads to increased T_REG_ abundance (Figure 1). Thus, despite observing increased PD-1 expression, our current data do not allow us to determine whether ITK knockdown enhances T_REG_ suppression on a per-cell basis. Nonetheless, we can conclude that T_REGs_ differentiated *in vitro* from naïve human CD4+ T cells are bona fide regulatory T cells and that ITK knockdown can promote increased differentiation (Figure 1) of functionally suppressive (Figure 4) human Foxp3+ T_REGS_.

### A pharmacological ITK/RLK kinase inhibitor abrogates the effect of ITK knockdown on T_REG_ differentiation

Studies in ITK deficient mice have fostered significant interest in targeting this kinase in autoimmune and inflammatory diseases [6, 7, 9, 18]. The novel covalent inhibitor PRN694 targets mainly ITK and RLK, and it blocks T cell activation, proliferation, and T helper cell differentiation [2, 3].

Furthermore, this compound ameliorated colitis progression in the T cell transfer colitis model by blunting both Th1 and Th17 responses [2]. However, in contrast to ITK-deficient mice, PRN694-treated mice exhibit reduced T_REG_ numbers, suggesting that RLK may play a distinct role in T_REG_ differentiation [2, 6]. In agreement with the findings in mice, PRN694 inhibited human T cell activation and T_REG_ differentiation in a dose-dependent manner, thus abrogating the increase in T_REGS_ observed upon ITK knockdown (Figure 5).

**Figure 5.**
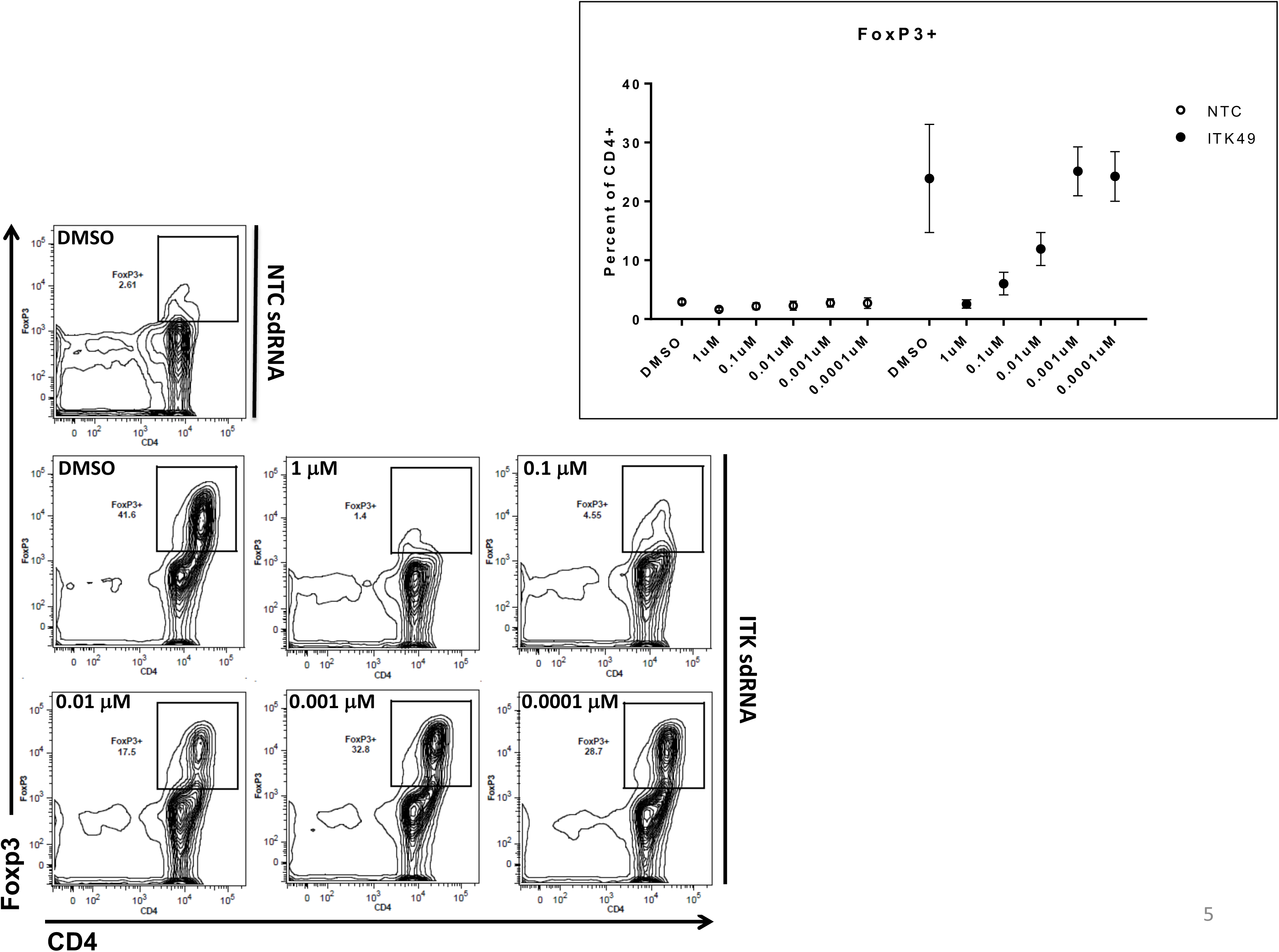
Human naïve CD4 T cells were treated with non-targeting (NTC) or ITK targeting sdRNA followed by the ITK/RLK kinase inhibitor PRN694 or DMSO control. Cells were then activated under T_REG_ polarizing conditions (IL-2, TGFβ). After four days, the CD4+ T cells were analyzed for Foxp3 expression by flow cytometry. N=3 donors.

Since RLK expression is suppressed after T cell activation [15], this Tec kinase is likely to be involved in early T cell activation events which are required to drive both T helper and T_REG_ differentiation. We attempted to isolate the effect of RLK on T_REG_ differentiation using sdRNA knockdown, but none of the eleven sdRNA constructs we tested alone or in combination reduced Txk gene expression (data not shown). Thus, while it is possible that RLK and ITK play distinct roles in T_REG_ and T helper differentiation, at this point we cannot determine whether these Tec kinases play redundant or reciprocal roles in this regard.

## Discussion

The TEC family of tyrosine kinases, including ITK and RLK, are critical for TCR-mediated T cell activation and T helper cell differentiation, and they are likely to play distinct roles in these processes. We show here for the first time that ITK is a negative regulator of human Foxp3+ T_REG_ and Th1 development, with a reciprocal effect on Th17 differentiation. Our findings extend recent observations in ITK deficient mice, which display enhanced T_REG_ development *in vivo* and *in vitro* and diminished T effector development and function [6, 7, 9].

In addition, we observed a marked increase in IFNγ production by Th1 polarized cells after ITK knockdown. ITK has a well-established role in Th2 mediated inflammation, but reports are conflicting on its role in Th1 effector function [5, 7, 9, 19-21]. While some data indicate that ITK positively regulates Th1 development and effector function in mouse T cells [7, 9, 20, 21], our data suggest that ITK is dispensable for Th1 effector function in primary human T cells, and that targeting ITK alone is therefore unlikely to be broadly immunosuppressive.

While RLK and ITK might fulfill some redundant roles [4], pharmacologic inhibition of both ITK and RLK in human naïve T cells globally blocked T cell activation and T_REG_ differentiation. This is consistent with observations in ITK/RLK double knockout mice, which showed markedly impaired TCR-mediated T cell proliferation [1]. Nonetheless, while ITK knockdown markedly enhanced human T_REG_ differentiation from naïve T cells, ITK knockdown followed by pharmacologic inhibition of ITK and RLK abrogated this effect, suggesting that, in contrast to ITK, RLK is not involved in T_REG_ suppression but is actually required for T_REG_ differentiation in the absence of ITK. ITK and RLK are both activated downstream of TCR ligation however, in contrast to ITK, RLK activation is independent of the PI3K-AKT axis [12] which is integral for the repression of FoxP3 protein and gene transcription [22, 23]. RLK also independently phosphorylates the adapter molecule SLP-76 leading to PLCγ activation [24], which explains its compensatory role in T cell activation and differentiation in the absence of ITK. Therefore, our data support a model whereby selective inhibition of ITK de-represses FoxP3 via inhibition of PI3K-AKT signaling while sparing RLK to promote PLCγ-mediated T cell activation and differentiation.

While we have shown that ITK knockdown enhances the *in vitro* differentiation of human T_REG_ with functional suppressive capacity *in vitro*, we cannot conclude whether ITK knockdown affects T_REG_ suppression on a per cell basis. In fact, even though initial ITK knockdown promotes T_REG_ differentiation, sdRNA content is expected to decrease due to initial T cell proliferation and/or degradation, and therefore our data cannot determine whether ITK is truly dispensable for T_REG_ suppressive function. The latter is a potentially important question, because a previous report showed that T_REG_ from ITK-deficient mice did not protect in a colitis model [8], although another paper showed that *ex vivo*-induced ITK-/- T_REG_ efficiently abrogated inflammation in a similar model [6].

The reciprocal effects of ITK on effector T cell and T_REG_ differentiation makes this a potentially attractive target in autoimmune and inflammatory diseases. However, the lack of small molecule inhibitors selective for ITK, sparing other Tec kinases such as RLK, makes this target technically challenging for drug development. Our current data in human T cells plus previous data in knockout mice [1] indicate that targeting both ITK and RLK might lead to excessive immunosuppression, abrogating the beneficial immunomodulatory effect of increasing T_REG_ differentiation that could be achieved by targeting ITK alone.

## Materials and Methods

### Cell culture

Naïve CD4+ CD45RA+ CD25-peripheral blood T cells were purchased from AllCells (Alameda, CA). T cells were cultured in RPMI 1640 media with pen/strep and 10% HI-FCS on wells coated with 1ug/ml of anti-CD3 (OKT3, eBioscience). Th cell polarization conditions were as follows. T_REGS_: 300U/ml of IL-2, 5ng/ml of rhTGFβ, 1ug/ml of anti-CD28. For Th17: IL-23 10ng/ml, TGFβ 0.5ng/ml, IL-1β 10ng/ml, IL-2 20U/ml, anti-INFγ 10ug/ml, anti-IL4 10ug/ml and anti-CD28 1ug/ml. For Th1: anti-IL4 10ug/ml, anti-CD28 1ug/ml, rh IL-2 20U/ml and rh IL-12 10ng/ml. For Th-null (Th0): 1ug/ml of anti-CD28 was added to the media. Cytokines were from R&D systems (Minneapolis, MN) and antibodies from eBioscience (San Diego, CA) unless otherwise noted.

### Knock down of ITK gene expression

Self-delivery siRNA (sdRNA) constructs were supplied by Advirna (Cambridge, MA). Cells were cultured in 1% serum in RPMI 1640 in non-activating conditions with sdRNA concentrations per manufacturer specifications for 24hrs, serum was added to 10% for an additional 24hrs then cells were transferred to media conditioned for T_REG_, Th17, Th1 or Th0 differentiation. Gene expression analysis was performed 72hr post-sdRNA addition to assess extent of ITK knockdown. Cells were analyzed after 4 days in T_REG_ or T_EFF_ polarizing conditions by flow cytometry for FoxP3+ regulatory T cells, CCR6 for Th17 and IFNγ for Th1. Several ITK-targeting sdRNA constructs were assessed in comparison to a non-targeting control (NTC) for ITK knock-down and FoxP3 upregulation and one construct (ITK49) was selected based on optimal knockdown efficiency and FoxP3+ T_REG_ induction (data not shown).

### *In vitro* suppression assay

T_REGS_ were differentiated as indicated above with ITK or NTC sdRNA for 4 days then collected and labeled with CellTrace Violet (Life Technologies). CD4+ peripheral blood responder cells (AllCells; Alameda, CA) were labeled with CFSE. T_REGS_ and responder cells were cultured with anti-CD3/CD28 activation beads (1:10 bead:responder cell ratio; Miltenyi) at T_REG_:responder cell ratios of 1:8 to 1:1 and were incubated together for 72 hours. After 72 hours, the cells were fixed and flow cytometry was run on the BD LSR-II to determine proliferation. Proliferation profile and division index analysis with FlowJo.

### qRT-PCR analysis

Total RNA was isolated from cultured T cells with the RNeasy 96 kit (Qiagen; Germantown, MD) and reverse transcribed into cDNA with qScript cDNA mastermix (Quanta Biosciences; Gaithersburg, MD). Quantitative RT-PCR (qPCR) was performed with Taqman primer-probe sets and Taqman universal mastermix (Applied Biosystesms). Data were collected on the Viaa7 Real-Time PCR system (Applied Biosystems™) and analyzed by the comparative Ct method (2– [delta][delta]Ct) with normalization to the mean Ct of endogenous control genes HPRT1, TBP and IPO8.

### Flow cytometry

Cells were stained for Foxp3 and other surface markers with the human regulatory t cell staining kit #1 (eBioscience). Cells were stained with the fixable Live/Dead aqua dye (Life Technologies) and CD4, CD25, Foxp3 and CCR6 antibodies. Prior to intracellular IFNγ staining, cells were stimulated with PMA/Ionomycin for 5 hours in the presence of Monensin for the last 4 hours of stimulation. Data were collected on the BD LSR-II and analysis was performed using FlowJo software.

### Luminex

Cytokines in culture media supernatants were measured with the Luminex platform using the Milliplex Human Cytokines 38 plex Immunology Assay (Millipore-Sigma; Billerica, Massachusetts).

### ITK/RLK compound

The dual ITK/RLK small molecule inhibitor was synthesized in-house (JNJ64461449) based on published data [2, 3]

### Statistics

Results are expressed as the mean +/- SEM. Statistical differences between 2 groups were calculated by unpaired student’s t test and among more than two groups by ANOVA. Graphs were generated with Prism-GraphPad software.

## References

1. Schaeffer, E.M., J. Debnath, G. Yap, D. McVicar, X.C. Liao, D.R. Littman, A. Sher, H.E. Varmus, M.J. Lenardo, and P.L. Schwartzberg, Requirement for Tec kinases Rlk and Itk in T cell receptor signaling and immunity. Science, 1999. 284(5414): p. 638–41.

2. Cho, H.S., H.M. Shin, H. Haberstock-Debic, Y. Xing, T.D. Owens, J.O. Funk, R.J. Hill, J.M. Bradshaw, and L.J. Berg, A Small Molecule Inhibitor of ITK and RLK Impairs Th1 Differentiation and Prevents Colitis Disease Progression. J Immunol, 2015. 195(10): p. 4822–31.

3. Zhong, Y., S. Dong, E. Strattan, L. Ren, J.P. Butchar, K. Thornton, A. Mishra, P. Porcu, J.M. Bradshaw, A. Bisconte, T.D. Owens, E. Verner, K.A. Brameld, J.O. Funk, R.J. Hill, A.J. Johnson, and J.A. Dubovsky, Targeting interleukin-2-inducible T-cell kinase (ITK) and resting lymphocyte kinase (RLK) using a novel covalent inhibitor PRN694. J Biol Chem, 2015. 290(10): p. 5960–78.

4. Sahu, N., A.M. Venegas, D. Jankovic, W. Mitzner, J. Gomez-Rodriguez, J.L. Cannons, C. Sommers, P. Love, A. Sher, P.L. Schwartzberg, and A. August, Selective expression rather than specific function of Txk and Itk regulate Th1 and Th2 responses. J Immunol, 2008. 181(9): p. 6125–31.

5. Sun, Y., I. Peng, J.D. Webster, E. Suto, J. Lesch, X. Wu, K. Senger, G. Francis, K. Barrett, J.L. Collier, J.D. Burch, M. Zhou, Y. Chen, C. Chan, J. Eastham-Anderson, H. Ngu, O. Li, T. Staton, C. Havnar, A. Jaochico, J. Jackman, S. Jeet, L. Riol-Blanco, L.C. Wu, D.F. Choy, J.R. Arron, B.S. McKenzie, N. Ghilardi, M.H. Ismaili, Z. Pei, J. DeVoss, C.D. Austin, W.P. Lee, and A.A. Zarrin, Inhibition of the kinase ITK in a mouse model of asthma reduces cell death and fails to inhibit the inflammatory response. Sci Signal, 2015. 8(405): p. ra122.

6. Gomez-Rodriguez, J., E.A. Wohlfert, R. Handon, F. Meylan, J.Z. Wu, S.M. Anderson, M.R. Kirby, Y. Belkaid, and P.L. Schwartzberg, Itk-mediated integration of T cell receptor and cytokine signaling regulates the balance between Th17 and regulatory T cells. J Exp Med, 2014. 211(3): p. 529–43.

7. Kannan, A., Y. Lee, Q. Qi, W. Huang, A.R. Jeong, S. Ohnigian, and A. August, Allele-sensitive mutant, Itkas, reveals that Itk kinase activity is required for Th1, Th2, Th17, and iNKT-cell cytokine production. Eur J Immunol, 2015. 45(8): p. 2276–85.

8. Huang, W., A.R. Jeong, A.K. Kannan, L. Huang, and A. August, IL-2-inducible T cell kinase tunes T regulatory cell development and is required for suppressive function. J Immunol, 2014. 193(5): p. 2267–72.

9. Deakin, A., G. Duddy, S. Wilson, S. Harrison, J. Latcham, M. Fulleylove, S. Fung, J. Smith, M. Pedrick, T. McKevitt, L. Felton, J. Morley, D. Quint, D. Fattah, B. Hayes, J. Gough, and R. Solari, Characterisation of a K390R ITK kinase dead transgenic mouse--implications for ITK as a therapeutic target. PLoS One, 2014. 9(9): p. e107490.

10. Berg, L.J., L.D. Finkelstein, J.A. Lucas, and P.L. Schwartzberg, Tec family kinases in T lymphocyte development and function. Annu Rev Immunol, 2005. 23: p. 549–600.

11. Labno, C.M., C.M. Lewis, D. You, D.W. Leung, A. Takesono, N. Kamberos, A. Seth, L.D. Finkelstein, M.K. Rosen, P.L. Schwartzberg, and J.K. Burkhardt, Itk functions to control actin polymerization at the immune synapse through localized activation of Cdc42 and WASP. Curr Biol, 2003. 13(18): p. 1619–24.

12. Debnath, J., M. Chamorro, M.J. Czar, E.M. Schaeffer, M.J. Lenardo, H.E. Varmus, and P.L. Schwartzberg, rlk/TXK encodes two forms of a novel cysteine string tyrosine kinase activated by Src family kinases. Mol Cell Biol, 1999. 19(2): p. 1498–507.

13. Czar, M.J., J. Debnath, E.M. Schaeffer, C.M. Lewis, and P.L. Schwartzberg, Biochemical and genetic analyses of the Tec kinases Itk and Rlk/Txk. Biochem Soc Trans, 2001. 29(Pt 6): p. 863–7.

14. Wang, X., S.E. Boyken, J. Hu, X. Xu, R.P. Rimer, M.A. Shea, A.S. Shaw, A.H. Andreotti, and Y.H. Huang, Calmodulin and PI(3,4,5)P(3) cooperatively bind to the Itk pleckstrin homology domain to promote efficient calcium signaling and IL-17A production. Sci Signal, 2014. 7(337): p. ra74.

15. Sommers, C.L., K. Huang, E.W. Shores, A. Grinberg, D.A. Charlick, C.A. Kozak, and P.E. Love, Murine txk: a protein tyrosine kinase gene regulated by T cell activation. Oncogene, 1995. 11(2): p. 245–51.

16. Fowell, D.J., K. Shinkai, X.C. Liao, A.M. Beebe, R.L. Coffman, D.R. Littman, and R.M. Locksley, Impaired NFATc translocation and failure of Th2 development in Itk-deficient CD4+ T cells. Immunity, 1999. 11(4): p. 399–409.

17. Schaeffer, E.M., G.S. Yap, C.M. Lewis, M.J. Czar, D.W. McVicar, A.W. Cheever, A. Sher, and P.L. Schwartzberg, Mutation of Tec family kinases alters T helper cell differentiation. Nat Immunol, 2001. 2(12): p. 1183–8.

18. Ferrara, T.J., C. Mueller, N. Sahu, A. Ben-Jebria, and A. August, Reduced airway hyperresponsiveness and tracheal responses during allergic asthma in mice lacking tyrosine kinase inducible T-cell kinase. J Allergy Clin Immunol, 2006. 117(4): p. 780–6.

19. Gomez-Rodriguez, J., N. Sahu, R. Handon, T.S. Davidson, S.M. Anderson, M.R. Kirby, A. August, and P.L. Schwartzberg, Differential expression of interleukin-17A and -17F is coupled to T cell receptor signaling via inducible T cell kinase. Immunity, 2009. 31(4): p. 587–97.

20. Kannan, A.K., D.G. Kim, A. August, and M.S. Bynoe, Itk signals promote neuroinflammation by regulating CD4+ T-cell activation and trafficking. J Neurosci, 2015. 35(1): p. 221–33.

21. Kannan, A.K., N. Sahu, S. Mohanan, S. Mohinta, and A. August, IL-2-inducible T-cell kinase modulates TH2-mediated allergic airway inflammation by suppressing IFN-gamma in naive CD4+ T cells. J Allergy Clin Immunol, 2013. 132(4): p. 811–20 e1-5.

22. Merkenschlager, M. and H. von Boehmer, PI3 kinase signalling blocks Foxp3 expression by sequestering Foxo factors. J Exp Med, 2010. 207(7): p. 1347–50.

23. Dang, E.V., J. Barbi, H.Y. Yang, D. Jinasena, H. Yu, Y. Zheng, Z. Bordman, J. Fu, Y. Kim, H.R. Yen, W. Luo, K. Zeller, L. Shimoda, S.L. Topalian, G.L. Semenza, C.V. Dang, D.M. Pardoll, and F. Pan, Control of T(H)17/T(reg) balance by hypoxia-inducible factor 1. Cell, 2011. 146(5): p. 772–84.

24. Schneider, H., B. Guerette, C. Guntermann, and C.E. Rudd, Resting lymphocyte kinase (Rlk/Txk) targets lymphoid adaptor SLP-76 in the cooperative activation of interleukin-2 transcription in T-cells. J Biol Chem, 2000. 275(6): p. 3835–40.

